# Detection of Feline Coronavirus RNA in Cats with Feline Infectious Peritonitis and their Housemates

**DOI:** 10.1101/2025.05.29.656645

**Authors:** Phoenix M. Shepherd, Amy Elbe, Brianna Lynch, Erin Lashnits, Robert Kirchdoerfer

## Abstract

Feline coronavirus (FCoV), the causative agent behind feline infectious peritonitis (FIP), is one of the biggest infectious threats to feline health. Despite this threat, the tissue distribution and viral RNA levels in cats infected with feline coronaviruses are poorly understood in the context of natural infection. Here, we used quantitative reverse-transcription PCR (qRT-PCR) to examine viral RNA levels from different sampling sites in both cats that have been clinically suspected of FIP and their feline housemates. We show that the distribution and amount of FCoV viral RNA does not differ between infected FIP cats and their feline housemates in blood, conjunctiva, or feces. Furthermore, in all FIP and non-FIP cases, viral RNA levels were higher in fecal samples than the blood. Taken together these results indicate a need for closer examination of FCoVs and how they cause FIP.

---

Coronaviruses are known for their ability to infect and subsequently cause disease in a wide variety of host species. Several human coronaviruses, such as MERS-CoV, SARS-CoV, and SARS-CoV-2, emerged due to spillover events from animal to human populations [1-5], likely because of promiscuous receptor use and due to the high rate at which these viruses mutate [6-8]. Feline coronaviruses are members of the *Alphacoronavirus* genus (family *Coronaviridae*, order *Nidovirales*) and are related to canine coronaviruses (CCoV) infecting dogs as well as transmissible gastroenteritis and porcine respiratory coronavirus (PRCV) virus infecting pigs [9]. Like their canine counterparts, FCoVs circulate in two serotypes. Type I is the most common serotype in the United States, accounting for as much as 80% of natural infections [10-11]. While less common, type II feline coronaviruses also circulate in the US [10-13]. FCoVs have remained an enigma in the scientific community due to their ability to cause both mild and severe diseases, an ability that is still poorly understood, yet is of paramount importance in developing antiviral and vaccination strategies.

FCoVs are classified into two pathotypes by the type of disease they cause. In most cases, the virus is entirely asymptomatic or causes mild gastrointestinal disease characterized by diarrhea or appetite loss [14-16]. In these cases, the causative virus is referred to as feline enteric coronavirus (FECV). Through unknown mechanisms, FCoVs may also cause a severe systemic disease known as feline infectious peritonitis (FIP).

The viral pathotype is then termed feline infectious peritonitis virus (FIPV). It is hypothesized that FIPV is not transmissible from cat to cat, but rather is the result of a parental FECV mutating over the course of infection to develop into FIPV and cause FIP. FIP can manifest in many forms, some of which are neurotropic [17-20]. FIP is most commonly recognized by the presence of granulomatous lesions and inflammation in several organs, such as the liver, kidney, and spleen. In some cases, fibrinous, granulomatous serositis [18-21] occurs, which results in the formation of protein-rich effusions in body cavities of diseased cats and is termed as “wet” FIP [19-20]. Cats afflicted with neurotropic “dry” FIP may experience seizures, ocular disease, behavioral changes, ataxia, and more [18-20]. Recently, an alarming outbreak of FIPV in Cyprus has sparked scientific interest in understanding how FCoVs evolve into new pathotypes within hosts to cause severe disease [21].

The gold standard for detection of FCoVs is to use *in situ* hybridization probes targeting the FCoV nucleocapsid or matrix protein in fixed tissues from biopsies or necropsies [18-19]. Treatment options for FIP are limited. Recently, nucleoside analogs GS-441524 and the related antiviral remdesivir, used to treat SARS-CoV-2 infections, have both been shown to be effective for curing cats of FIP with limited side effects [22-25]. In addition, an inhibitor of the FCoV viral protease, GC376, has also been shown as effective against viral replication in cats, albeit to a lesser extent [26-28]. While effective, these treatments are not preventative and tend to be expensive. Greater insight into the prevalence of feline coronavirus across different sites within the cat are needed to develop better vaccination and treatment strategies. Progress is hindered by the lack of reliable methodologies to measure feline coronaviruses in a variety of samples as well as a lack of matched controls between FIPV- and FECV-infected cats. Here, we have developed a Taqman-probe based qRT-PCR assay that targets the 3’ end of the feline coronavirus RNA genome to quantify viral RNA in both cats infected with FIPV and FECV. We have recruited a cohort of cats who have been clinically suspected of FIP or are housemates to those FIP cats, with the hypothesis that both groups may have experienced infection of the same parental virus. We use qRT-PCR in both symptomatic FIP cats and their housemates to better understand viral spread in both cases of severe FIP and asymptomatic FCoV infections.

## Assessing specificity of the qRT-PCR assay

To examine how specific our assay was at detecting feline coronaviruses, we designed additional standard curves, based on homologous sequences in two closely related alphacoronaviruses: canine coronavirus (CCoV) and porcine respiratory coronavirus (PRCV). These results are shown in Figure S1. There was sufficient cross-reactivity between our FCoV primers, CCoV, and to a lesser extent PRCV, so we cannot rule out the cats presented in this study were not infected with PRCV or CCoV rather than FCoV; however, to our knowledge, there is no biochemical evidence that these viruses can replicate in feline hosts.

Next, we wanted to know if our assay is sensitive to background RNA or DNA from feline hosts. We first isolated RNA from two different feline cell lines: Fcwf-4 macrophages (ATCC CRL-278) and AK-D lung epithelium (ATCC CCL-150). After qRT-PCR, both samples were below the limit of detection (data not shown). We also recruited nine healthy housecats with unknown infection histories from single-cat households and sampled them using the methodology described above. Data displayed in Figure S 1 shows some quantifiable viral RNA in one or two cats from each sample (other than plasma, which had undetectable levels in all cats tested). This level of positivity is lower than what has been previously reported in healthy housecats [31-32].

## Sites and Levels of FCoV RNA do not Distinguish Clinical FIP

### Plasma

To understand the systemic spread of FCoV in both cats with and without FIP, we used our qRT-PCR assay to determine the total amount of viral RNA in cat plasma (Figure 1A). Results show that overall positivity in both groups was low (Table 1), meaning most cats tested below the limit of detection in our assay. Those cats that did test above the limit of quantification tended to have low amounts of viral RNA in their plasma. When comparing diseased and non-diseased cats, we observed no significant difference in viral RNA levels between positive cats from the two groups (Figure 1A, P = 0.2134).

**Table 1.** Metadata from our cohort displayed percent positivity in all cats. Data was sorted into quantifiable, detected, or below the limit of detection.

|  | **Plasma** | **Whole**<br>**Blood** | **Fecal**<br>**Swab** | **Conjunctival**<br>**Swab** | **Effusion** |
| --- | --- | --- | --- | --- | --- |
| % Quantifiable | 22.1% | 35.0% | 72.6% | 67.9% | 75.0% |
| % Detected | 19.1% | 15.0% | 8.1% | 10.7% | 0.0% |
| % Below the<br>LoD | 44.1% | 25.0% | 9.7% | 7.1% | 25.0% |
| % Undetected | 14.7% | 25.0% | 9.7% | 14.3% | 0.0% |

**Figure 1.**
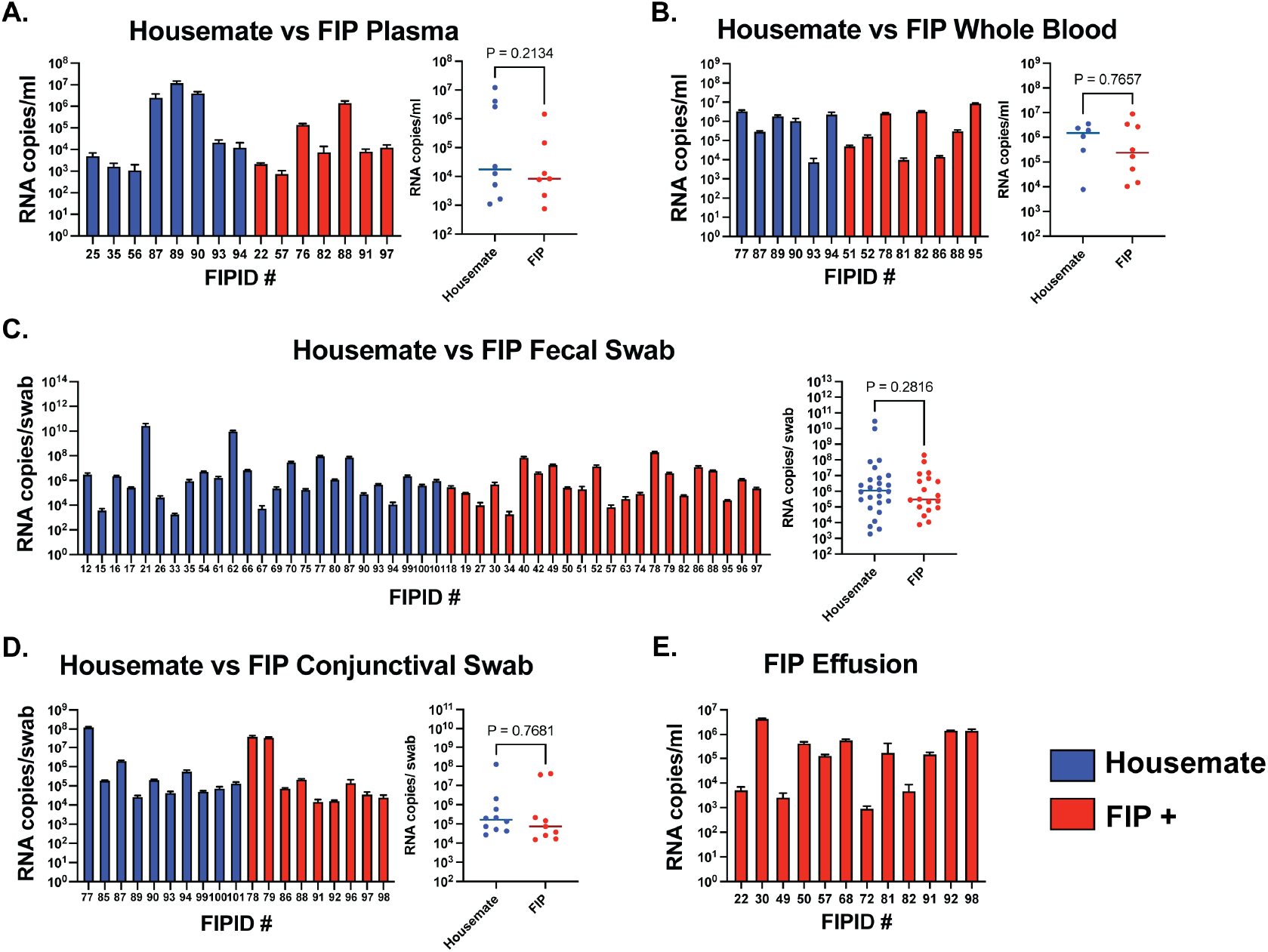
Quantifiable qRT-PCR data as RNA copies/ml or RNA copies/swab in each sample type: A) plasma; B) whole blood; C) fecal swabs; D) conjunctival swabs; and E) effusion. Samples that were detected but not quantifiable or were below the limit of detection are not shown. Unpaired t-test results are displayed as P-values. Statistical significance was defined as P < 0.05.

### Whole Blood

Some studies have found [32-36] that FCoV RNA is cell associated, meaning acellular plasma may result in negative or low qRT-PCR signals. We collected whole blood samples prior to centrifugation for qRT-PCR. Aggregated results show the same prevalence of positivity in both FIP and non-FIP cats. However, average RNA levels in whole blood were slightly increased compared to plasma, although the difference is not significant (Figure 1B). When comparing RNA levels in whole blood samples of FIP and non-FIP cats, there was also no significant difference between the two groups, similar to the results for plasma samples (Figure 1B, P = 0.7657).

### Fecal Swabs

To measure the amount of enteric FCoV RNA, we performed qRT-PCR on fecal swab samples. In contrast to the plasma and whole blood samples, we observed high amounts of viral RNA extracted from fecal swabs (Figure 1C). Additionally, both FIP and non-FIP cat groups presented greater overall positivity (Table 1). We observed comparable amounts of viral RNA between FIP and non-FIP cats (Figure 1C, P = 0.2816). The percent of positivity and high RNA levels in both groups, leads us to speculate that there may be viral shedding in both FIP cats and their feline housemates.

### Conjunctival Swabs

Because some forms of feline coronaviruses can manifest as ocular disease [18, 21], we sampled a portion of our cats for the presence of feline coronavirus in the conjunctiva using the same methodology as our fecal swabs.

Surprisingly, most cats showed detectable levels of viral RNA in these samples (Table 1). Similar to other sample sites, there was no statistically significant difference between cats with and without FIP disease (Figure 1D, P = 0.7681).

### Effusion

We also used our qRT-PCR assay to measure the level of viral RNA in effusions from “wet” FIP cats. Here, we observed high levels of viral RNA as well as a large percentage of overall positivity (Figure 1E, Table 1). The amounts of viral RNA we observed here are similar to those reported previously [33-36, 38].

## Viral RNA levels are not different between FIP cats and their housemates

Due to the lack of significant differences in FIP cats and their housemates in the overall cohort, we decided to look more closely at individual households of cats, either in pairs or trios. We sampled ten multi-cat households from our dataset and compared viral RNA levels among members of that household (Figure 2). Results from these comparisons agree with our data from the entire study cohort. Although there are significant differences in individual sample types, in some groups, the FIP cats had significantly higher viral RNA, while in others, the housemates had higher levels of viral RNA (Figure 2). Given the unknown infection history of sampled cats, we can make no conclusions as to the timing or progression of FCoV infections in individual households.

**Figure 2.**
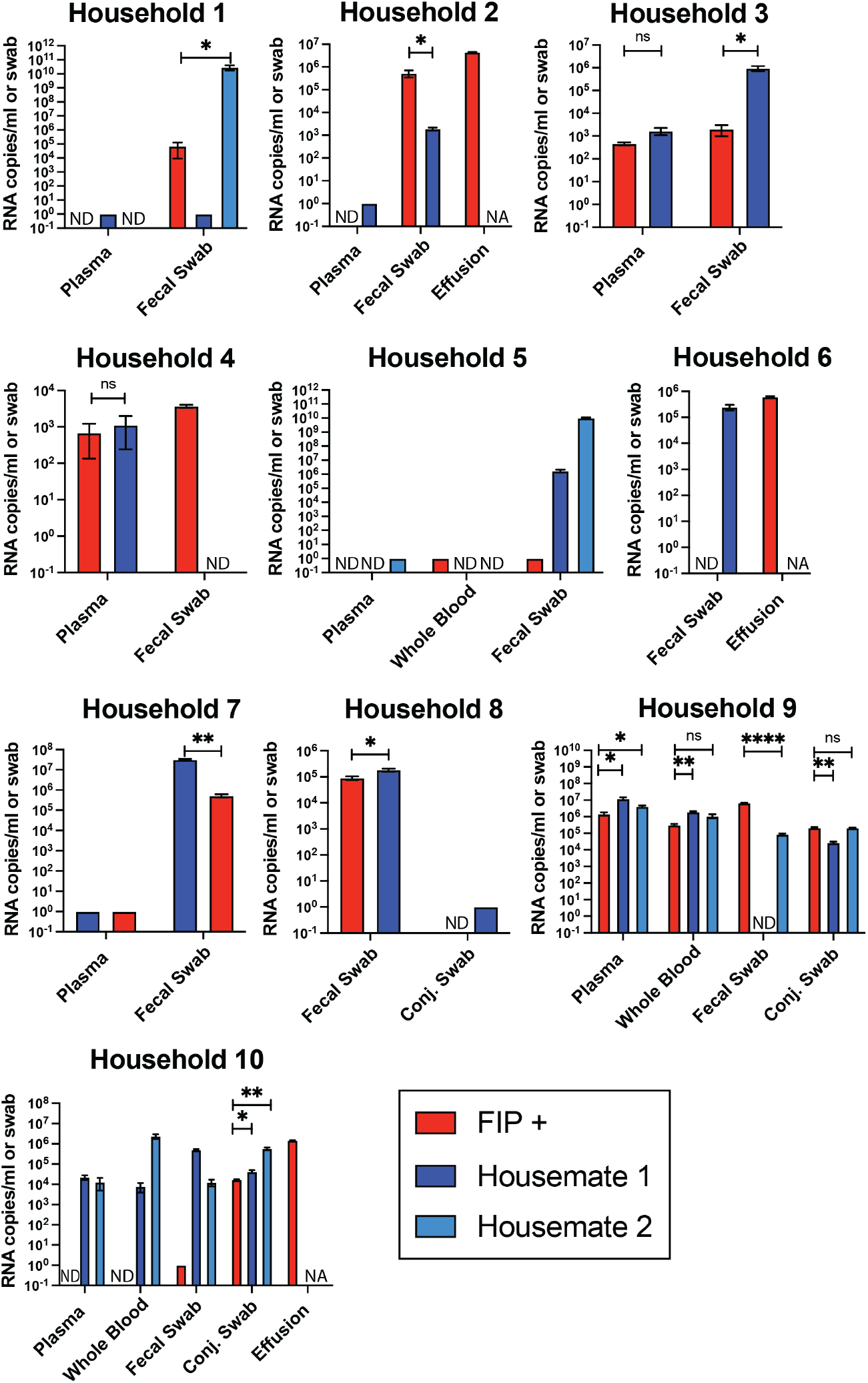
Comparisons between FIP cats and their housemates in selected households. Households selected based on qRT-PCR results. Samples detectable but not quantifiable were given a value of 1. Samples below the limit of detection were given a value of 0 and labeled as no data. NA, not applicable; ND, no data; ns, not significant; * = P < 0.05; ** = P < 0.01; *** = P < 0.001; **** = P < 0.0001 based on paired t-tests.

## Discussion

This study shows that both cats clinically suspected of FIP as well as their housemates had detectable levels of FCoV RNA in various sample types. The low-to-moderate levels of RNA observed in the blood (both plasma and whole blood) as well as the higher amounts of FCoV RNA isolated from fecal swabs, conjunctival swabs, and effusions agree with past studies showing varying levels of viral RNA in cats with and without clinical FIP [16, 33-38]. Additionally, our study uses the TaqMan FAM/ZEN/IABQ probe based qRT-PCR system to target a conserved sequence between primer binding sites. While single-quenched probes have been developed [38, 40-41], to our knowledge, a dual-quenched probe has not been used before to study clinical samples of feline coronaviruses. Dual-quenched probes have been shown to offer increased sensitivity in other qPCR assays [42-44].

The assays here use a standard curve based on absolute quantification of RNA copies. Such methods allow the quantification and comparison of different sample types across different cats. However, sampling from client owned cats, we acknowledge that the timing of viral infection differs between cats sampled in this study. All time points used in this present study are when the FIP cats presented to the clinic. Those cats who are very early or much later in the course of viral infection may have had undetectable levels of virus at some sampling sites due to virus clearance or low viral replication.

In our cohort of cats, the levels of viral RNA did not differ significantly between cats who were suspected of FIP and their housemates, indicating that detecting viral RNA is not enough to conclusively diagnose FIP. In unexposed, healthy cats with unknown infection histories, we observed a much lower percentage of quantifiable RNA in all samples compared to the feline housemate group (Table 1, Table S1). The qRT-PCR assay presented here, like all viral qPCRs, does not differentiate between actively replicating virus and residual viral RNA. However, we do note that we were able to detect FCoV RNA in the blood (whole blood and plasma) of cats who did not have FIP. Such findings lead us to confirm that systemic spread of FCoV alone is not sufficient to cause FIP, agreeing with several other studies [37-40]. Also, we were able to detect high levels of viral RNA in the feces of both FIP cats and their housemates. Generally, we quantified higher amounts of feline coronavirus RNA derived from the feces than other sample types. We speculate that the presence of FCoV RNA in these fecal samples may suggest that shedding of FCoV is occurring in both FIP cats and their housemates. This study did not assess the viral populations that may be shed in the feces.

Cats were recruited with the hypothesis that infections in multi-cat households stem from a single founder virus which developed into FIP in some cats but not in their housemates. The high potential for relatedness among infecting viruses within a household is a controlling factor allowing finer comparisons between FIP and non-FIP pathotype viruses. In this case study, we note the same trends among cats from the same household as in the aggregated cohort of cats. However, there are no correlations in viral RNA levels with FIP disease when examining individual multi-cat households.

We conclude that the development of FIP does not correlate with the amount, nor the anatomical location of viral RNA. Thus, the exact reasons for the emergence of FIP within cats remains unknown. However, the methodologies presented in this case study illustrate a new way of looking at feline coronavirus infections in the context of natural infection by directly comparing different sample types from two cohorts of cats: those with clinically suspected FIP and their housemates. Additional analyses using viral genome sequencing may shed light about the emergence of this virus and the viral populations that arise before and after FIP diagnosis.

## Materials and Methods

### Recruitment of study cats and housemates

Cats were enrolled at the University of Wisconsin-Madison Veterinary Teaching Hospital, University of Wisconsin Veterinary Care after evaluation of symptoms based on owner report and veterinary review. Each cat was assigned an FIP Identification (FIPID) number to track sample intake and household relations. Cats were diagnosed according to the 2022 AAFP/EveryCat FIP diagnosis guidelines and the European Advisory Board on Cat Diseases FIP guidelines [29]. Definitive diagnosis of FIP based on immunohistochemistry was not required for inclusion, rather cats could be included based on clinical suspicion. Cats with a confirmed alternative diagnosis were excluded.

Cats were not included if an alternative diagnosis (such as toxoplasmosis, congestive heart failure, or septic pleuritis) was reached. Cats that had already begun treatment with antiviral agents were excluded from the analysis.

Blood, fecal and conjunctival swabs, and effusion (if present) from cases of probable FIP were taken and owners were asked if the cat had healthy feline housemates. Housemates were recruited at a later date. Nine unexposed, healthy cats from single-cat households were also recruited to assess specificity of our assay. Due to the nature of the study, infection histories of all cats used in this case study are unknown. All cats used in this study were enrolled with the owners’ consent and had IACUC approval protocol #V006485. Some cats received clinical treatment before sampling and were not included in analysis

### Feline sample processing

#### Blood draws

2-3 mL of whole blood, depending on cat weight and anemic status, were drawn from cats using a vacutainer blood draw system (*BDVacutainier Tubes 367856 and 387812)*. Blood was drawn into 3 mL vacutainer tubes spray-coated with 5.4 mg of K_2_EDTA (whole blood). Whole blood from the K_2_EDTA coated tubes was aliquoted prior to centrifugation at 1,400 x g for 15 minutes. The lower density layer consisting of either plasma was removed and spun again at 500 x g for eight minutes to pellet any remaining cells. Whole blood and plasma were then aliquoted into sterile 1.5 mL Eppendorf tubes and frozen at -80°C.

#### Effusion

Total effusion volume (between 100 µL and 10 mL, depending on availability) was collected in a sterile 1.5 mL Eppendorf tube(s). Effusion was spun at 500 x g for eight minutes and the supernatant was aliquoted into sterile 1.5 mL Eppendorf tubes and frozen at -80°C.

#### Fecal Swabs

Anal swabs *(Copan FLOQSwabs 220250*) were used to collect fecal samples. Swabs were inserted dry and swirled on the swab axis for 10 s to collect feces. Fecal swabs were then placed into 500 µL of 1X DNA/RNA Shield *(Zymo Research R1200)* and either frozen at -80°C or immediately processed to collect RNA. Swabs were removed via sterilized forceps and wrung out on sides of the tubes. The 1X DNA/RNA Shield tube containing wrung-out fecal samples were relabeled with appropriate FIPIDs and frozen at -80°C.

#### Conjunctival Swabs

Conjunctival swabs *(Copan FLOQSwabs 220250*) were used to collect ocular samples. Cat conjunctiva was swabbed for 10 s and placed into 1X DNA/RNA Shield *(Zymo Research R1200)* and processed as described with fecal swabs above.

### Extraction of feline coronavirus RNA and synthesis of cDNA

100 uL of whole blood, plasma, effusion, or fecal swab 1X DNA/RNA Shield was used for viral RNA extraction. Extraction was performed by using Viral Quick RNA Extraction Kit *(Zymo Research R1034)* and RNA Cleanup Kit *(Zymo Research R1019)* following the protocols for DNase treatment. Prior to adding the Viral RNA Buffer, 2 µL of baker’s yeast (1 mg/mL) RNA *(Millipore Sigma R6750)* was added to the sample as carrier RNA. RNA was immediately used in reverse-transcription or stored at -80°C. For FCoV cDNA synthesis, 10 µL of RNA was added to 6 µL of nuclease-free water and 4 uL of Maxima H-minus RT master mix *(ThermoFisher EP0752)*. Thermocycler conditions were 25°C for 10 minutes, 65°C for 15 minutes, 85°C for 5 minutes, 10°C hold. cDNA was stored at -20°C until used in the qRT-PCR assay.

### Taqman-probe qRT-PCR

#### In vitro Transcription (IVT) of RNA Standards

A 130-nucleotide sequence matching the conserved 3’ end of the FCoV genome was synthesized as DNA in a pBluescript vector, containing a 5’ T7 promoter and 3’ restriction sites *(GenScript)*. Plasmid DNA was linearized via restriction digestion using the *Xbal* restriction enzyme and IVT was conducted using NEB IVT buffer and T7 RNA polymerase *(New England Biolabs M0251)*. Standards were then treated with DNaseI *(New England Biolabs M0303)* and an RNA Cleanup Kit *(Zymo Research R1019)*. RNA was quantified by UV absorbance *(DeNovix Inc)* and diluted for a standard curve. Standard curves were formed by serially diluting RNA stocks from 10^8^ copies of RNA/mL to 10^0^ copies of RNA/mL and reverse transcribed as described above.

#### Primers and Probes

To locate target sites to accommodate primer binding, we used ClustalOmega [30] software to align the genomes of 71 publicly available FCoV genomes. Primers and probes targeting a conserved region (accession number GQ152141.1 nucleotide position 28,646-28,775) in FCoV genomes were ordered through Integrated DNA Technologies (IDT). Forward sequence is CAACCCGATGTTTAAAACTGGT while the reverse sequence is CTAAATCTAGCATTGCCAAATCAAAT. The TaqMan probe forward sequence was manufactured by IDT and contains a 5’ 6-FAM fluorophore, a ZEN quencher, and a 3’ Iowa Black fluorescein quencher. The probe sequence is /56-FAM/CTACTCTTG/ZEN/TACAGAATGGTAAGCACGT/3IABkFQ/.

#### Quantitative PCR

2 µL of sample cDNA or standard was added to a clear plastic 96 well plate *(GeneMate, 490003-822)*. Mastermix containing forward, reverse and probe oligonucleotides (final concentration 1 µM), 2X iTaq Universal Probes Supermix *(BioRad, 1725130)*, and nuclease free water was added to each well and mixed. Total reaction volume was 12.5 µL. A BioRad CFX qPCR machine was used to read FAM fluorescence. RNA copies/ml were calculated based on threshold count in each sample. All samples and standards were run in triplicate. Assays were accepted if PCR efficiency was between 95%-110% with a standard curve R-squared value of >0.99. Limits of quantification (LoQ) and limits of detection (LoD) were determined by the BioRad CFX qPCR software, using 10% of the maximum relative fluorescence unit (RFU) of the least-dilute standard as a threshold for amplification. Each sample was categorized as quantifiable, detectable, below the limit of detection, or negative based on the number of cycles used to cross the threshold and the average RFU of the last five cycles. LoQ and LoD were based on the performance of the standards per assay. For these assays, the LoQ was typically between 700-1000 RNA copies/mL.

#### Statistics

Statistical analysis was performed using GraphPad Prism version 10.2.1 for MacOS (GraphPad Software, Boston, Massachusetts USA). To compare between unmatched FIP cats and their housemates, we used unpaired student *t-tests* and defined statistical significance by a P-value of less than 0.05. To compare samples from matched cats of the same household, a paired *t-test* was used. All samples were run as technical triplicates, including standards. Each technical triplicate was then averaged and used in statistical analysis. Only quantifiable samples were used in statistical analysis.

## References

1. Cui J, Li F, Shi ZL. Origin and evolution of pathogenic coronaviruses. Nat Rev Microbiol. 2019 Mar;17(3):181–192. doi: 10.1038/s41579-018-0118-9. PMID: 30531947; PMCID: PMC7097006.

2. Woolhouse ME, Haydon DT, Antia R. Emerging pathogens: the epidemiology and evolution of species jumps. Trends Ecol Evol. 2005 May;20(5):238–44. doi: 10.1016/j.tree.2005.02.009. PMID: 16701375; PMCID: PMC7119200.

3. Widagdo W, Okba NMA, Stalin Raj V, Haagmans BL. MERS-coronavirus: From discovery to intervention. One Health. 2016 Dec 23;3:11–16. doi: 10.1016/j.onehlt.2016.12.001. PMID: 28616497; PMCID: PMC5454172.

4. Yang Y, Liu C, Du L, Jiang S, Shi Z, Baric RS, Li F. Two mutations were critical for bat-to-human transmission of Middle East respiratory syndrome coronavirus. J Virol. 2015;89(17):9119–9123. doi: 10.1128/JVI.01279-15

5. Chan JF, To KK, Tse H, Jin DY, Yuen KY. Interspecies transmission and emergence of novel viruses: lessons from bats and birds. Trends Microbiol. 2013 Oct;21(10):544–55. doi: 10.1016/j.tim.2013.05.005. Epub 2013 Jun 14. PMID: 23770275; PMCID: PMC7126491.

6. Domingo E, García-Crespo C, Lobo-Vega R, Perales C. Mutation Rates, Mutation Frequencies, and Proofreading-Repair Activities in RNA Virus Genetics. Viruses. 2021 Sep 21;13(9):1882. doi: 10.3390/v13091882. PMID: 34578463; PMCID: PMC8473064.

7. Flores-Vega VR, Monroy-Molina JV, Jiménez-Hernández LE, Torres AG, Santos-Preciado JI, Rosales-Reyes R. SARS-CoV-2: Evolution and Emergence of New Viral Variants. Viruses. 2022 Mar 22;14(4):653. doi: 10.3390/v14040653. PMID: 35458383; PMCID: PMC9025907.

8. A Rohaim MF, El Naggar R M, Helal A M, Bayoumi M A, El-Saied M A, Ahmed K Z, Shabbir M, Munir M. Genetic Diversity and Phylodynamics of Avian Coronaviruses in Egyptian Wild Birds. Viruses. 2019 Jan 14;11(1):57. doi: 10.3390/v11010057. PMID: 30646528; PMCID: PMC6356246.

9. Chan JF, Lau SK, To KK, Cheng VC, Woo PC, Yuen KY. Middle East respiratory syndrome coronavirus: another zoonotic betacoronavirus causing SARS-like disease. Clin Microbiol Rev. 2015 Apr;28(2):465–522. doi: 10.1128/CMR.00102-14. PMID: 25810418; PMCID: PMC4402954.

10. Benetka V, Kübber-Heiss A, Kolodziejek J, Nowotny N, Hofmann-Parisot M, Möstl K. Prevalence of feline coronavirus types I and II in cats with histopathologically verified feline infectious peritonitis. Vet Microbiol. 2004 Mar 26;99(1):31–42. doi: 10.1016/j.vetmic.2003.07.010. PMID: 15019109; PMCID: PMC7117137.

11. Addie DD, Schaap IAT, Nicolson L, Jarrett O. Persistence and transmission of natural type I feline coronavirus infection. J Gen Virol. 2003 Oct;84(Pt 10):2735-2744. doi: 10.1099/vir.0.19129-0. PMID: 13679608.

12. Kummrow M, Meli ML, Haessig M, Goenczi E, Poland A, Pedersen NC, Hofmann-Lehmann R, Lutz H. Feline coronavirus serotypes 1 and 2: seroprevalence and association with disease in Switzerland. Clin Diagn Lab Immunol. 2005 Oct;12(10):1209–15. doi: 10.1128/CDLI.12.10.1209-1215.2005. PMID: 16210485; PMCID: PMC1247821.

13. Li C, Liu Q, Kong F, Guo D, Zhai J, Su M, Sun D. Circulation and genetic diversity of Feline coronavirus type I and II from clinically healthy and FIP-suspected cats in China. Transbound Emerg Dis. 2019 Mar;66(2):763–775. doi: 10.1111/tbed.13081. Epub 2018 Dec 5. PMID: 30468573; PMCID: PMC7168551.

14. Kipar A, Meli ML, Baptiste KE, Bowker LJ, Lutz H. Sites of feline coronavirus persistence in healthy cats. J Gen Virol. 2010 Jul;91(Pt 7):1698-707. doi: 10.1099/vir.0.020214-0. Epub 2010 Mar 17. PMID: 20237226.

15. Vogel L, Van der Lubben M, te Lintelo EG, Bekker CP, Geerts T, Schuijff LS, Grinwis GC, Egberink HF, Rottier PJ. Pathogenic characteristics of persistent feline enteric coronavirus infection in cats. Vet Res. 2010 Sep-Oct;41(5):71. doi: 10.1051/vetres/2010043. Epub 2010 Jul 23. PMID: 20663472; PMCID: PMC2939696.

16. Doenges SJ, Weber K, Dorsch R, Fux R, Fischer A, Matiasek LA, Matiasek K, Hartmann K. Detection of feline coronavirus in cerebrospinal fluid for diagnosis of feline infectious peritonitis in cats with and without neurological signs. J Feline Med Surg. 2016 Feb;18(2):104–9. doi: 10.1177/1098612x15574757. Epub 2015 Mar 3. PMID: 25736448.

17. Foley JE, Lapointe JM, Koblik P, Poland A, Pedersen NC. Diagnostic features of clinical neurologic feline infectious peritonitis. J Vet Intern Med. 1998 Nov-Dec;12(6):415–23. doi: 10.1111/j.1939-1676.1998.tb02144.x. PMID: 9857333; PMCID: PMC7167019

18. Stiles J. Ocular manifestations of feline viral diseases. Vet J. 2014 Aug;201(2):166–73. doi: 10.1016/j.tvjl.2013.11.018. Epub 2013 Dec 1. PMID: 24461645; PMCID: PMC7110540.

19. Kipar A, May H, Menger S, Weber M, Leukert W, Reinacher M. Morphologic features and development of granulomatous vasculitis in feline infectious peritonitis. Vet Pathol. 2005 May;42(3):321–30. doi: 10.1354/vp.42-3-321. PMID: 15872378.

20. Diaz JV, Poma R. Diagnosis and clinical signs of feline infectious peritonitis in the central nervous system. Can Vet J. 2009 Oct;50(10):1091-3. PMID: 20046611; PMCID: PMC2748294.

21. Attipa C., Warr AS., Epaminondas D., O’Shea M., Fletcher S., Malbon A., Lyraki M., Hammond R., Hardas A., Zanti A., Loukaidou S., Gentil M., Gunne-Moore D., Mazeri S., Tait-Burkard C. Emergence and spread of feline infectious peritonitis due to a highly pathogenic canine/feline recombinant coronavirus bioRxiv 2023.11.08.566182; doi: 10.1101/2023.11.08.566182

22. Krentz D, Zenger K, Alberer M, Felten S, Bergmann M, Dorsch R, Matiasek K, Kolberg L, Hofmann-Lehmann R, Meli ML, Spiri AM, Horak J, Weber S, Holicki CM, Groschup MH, Zablotski Y, Lescrinier E, Koletzko B, von Both U, Hartmann K. Curing Cats with Feline Infectious Peritonitis with an Oral Multi-Component Drug Containing GS-441524. Viruses. 2021 Nov 5;13(11):2228. doi: 10.3390/v13112228. PMID: 34835034; PMCID: PMC8621566.

23. Kokic G, Hillen HS, Tegunov D, Dienemann C, Seitz F, Schmitzova J, Farnung L, Siewert A, Höbartner C, Cramer P. Mechanism of SARS-CoV-2 polymerase stalling by remdesivir. Nat Commun. 2021 Jan 12;12(1):279. doi: 10.1038/s41467-020-20542-0. PMID: 33436624; PMCID: PMC7804290.

24. Green J, Syme H, Tayler S. Thirty-two cats with effusive or non-effusive feline infectious peritonitis treated with a combination of remdesivir and GS-441524. J Vet Intern Med. 2023 Sep-Oct;37(5):1784–1793. doi: 10.1111/jvim.16804. Epub 2023 Jul 4. PMID: 37403259; PMCID: PMC10472986.

25. Meli ML, Spiri AM, Zwicklbauer K, Krentz D, Felten S, Bergmann M, Dorsch R, Matiasek K, Alberer M, Kolberg L, von Both U, Hartmann K, Hofmann-Lehmann R. Fecal Feline Coronavirus RNA Shedding and Spike Gene Mutations in Cats with Feline Infectious Peritonitis Treated with GS-441524. Viruses. 2022 May 17;14(5):1069. doi: 10.3390/v14051069. PMID: 35632813; PMCID: PMC9147249.

26. Kim Y, Shivanna V, Narayanan S, Prior AM, Weerasekara S, Hua DH, Kankanamalage AC, Groutas WC, Chang KO. Broad-spectrum inhibitors against 3C-like proteases of feline coronaviruses and feline caliciviruses. J Virol. 2015 May;89(9):4942–50. doi: 10.1128/JVI.03688-14. Epub 2015 Feb 18. PMID: 25694593; PMCID: PMC4403489.

27. Pedersen NC, Kim Y, Liu H, Galasiti Kankanamalage AC, Eckstrand C, Groutas WC, Bannasch M, Meadows JM, Chang KO. Efficacy of a 3C-like protease inhibitor in treating various forms of acquired feline infectious peritonitis. J Feline Med Surg. 2018 Apr;20(4):378–392. doi: 10.1177/1098612x17729626. Epub 2017 Sep 13. PMID: 28901812; PMCID: PMC5871025.

28. Kim Y, Liu H, Galasiti Kankanamalage AC, Weerasekara S, Hua DH, Groutas WC, Chang KO, Pedersen NC. Reversal of the Progression of Fatal Coronavirus Infection in Cats by a Broad-Spectrum Coronavirus Protease Inhibitor. PLoS Pathog. 2016 Mar 30;12(3):e1005531. doi: 10.1371/journal.ppat.1005531. Erratum in: PLoS Pathog. 2016 May;12(5):e1005650. PMID: 27027316; PMCID: PMC4814111.

29. Thayer V, Gogolski S, Felten S, Hartmann K, Kennedy M, Olah GA. 2022 AAFP/EveryCat Feline Infectious Peritonitis Diagnosis Guidelines. J Feline Med Surg. 2022 Sep;24(9):905–933. doi: 10.1177/1098612x221118761. Erratum in: J Feline Med Surg. 2022 Dec;24(12):e676. doi: 10.1177/1098612×221126448. PMID: 36002137; PMCID: PMC10812230.

30. Madeira F., Pearce M., Tivey ARN., Basutkar P., Lee J., Edbali O., Madhusoodanan N., Kolesnikov A., Lopez R. (2022) Search and sequence analysis tools services from EMBL-EBI in 2022 Nucleic Acids Research, April 12, 2022; doi: 10.1093/nar/gkac240

31. Jähne S, Felten S, Bergmann M, Erber K, Matiasek K, Meli ML, Hofmann-Lehmann R, Hartmann K. Detection of Feline Coronavirus Variants in Cats without Feline Infectious Peritonitis. Viruses. 2022 Jul 29;14(8):1671. doi: 10.3390/v14081671. PMID: 36016293; PMCID: PMC9412601.

32. Can-Sahna K, Soydal Ataseven V, Pinar D, Oğuzoğlu TC. The detection of feline coronaviruses in blood samples from cats by mRNA RT-PCR. J Feline Med Surg. 2007 Oct;9(5):369–72. doi: 10.1016/j.jfms.2007.03.002. Epub 2007 May 2. PMID: 17478116; PMCID: PMC7128869.

33. Dong, B., Zhang, X., Zhong, X. et al. Prevalence of natural feline coronavirus infection in domestic cats in Fujian, China. Virol J 21, 2 (2024). 10.1186/s12985-023-02273

34. Pedersen NC, Eckstrand C, Liu H, Leutenegger C, Murphy B. Levels of feline infectious peritonitis virus in blood, effusions, and various tissues and the role of lymphopenia in disease outcome following experimental infection. Vet Microbiol. 2015 Feb 25;175(2-4):157-66. doi: 10.1016/j.vetmic.2014.10.025. Epub 2014 Nov 1. PMID: 25532961; PMCID: PMC7117444.

35. Doenges SJ, Weber K, Dorsch R, Fux R, Hartmann K. Comparison of real-time reverse transcriptase polymerase chain reaction of peripheral blood mononuclear cells, serum and cell-free body cavity effusion for the diagnosis of feline infectious peritonitis. J Feline Med Surg. 2017 Apr;19(4):344–350. doi: 10.1177/1098612x15625354. Epub 2016 Jul 9. PMID: 26787293.

36. Herrewegh AA, de Groot RJ, Cepica A, Egberink HF, Horzinek MC, Rottier PJ. Detection of feline coronavirus RNA in feces, tissues, and body fluids of naturally infected cats by reverse transcriptase PCR. J Clin Microbiol. 1995 Mar;33(3):684–9. doi: 10.1128/jcm.33.3.684-689.1995. PMID: 7751377; PMCID: PMC228014.

37. Fish EJ, Diniz PPV, Juan YC, Bossong F, Collisson EW, Drechsler Y, Kaltenboeck B. Cross-sectional quantitative RT-PCR study of feline coronavirus viremia and replication in peripheral blood of healthy shelter cats in Southern California. J Feline Med Surg. 2018 Apr;20(4):295–301. doi: 10.1177/1098612x17705227. Epub 2017 Apr 20. PMID: 28425327.

38. Lorusso E, Mari V, Losurdo M, Lanave G, Trotta A, Dowgier G, Colaianni ML, Zatelli A, Elia G, Buonavoglia D, Decaro N. Discrepancies between feline coronavirus antibody and nucleic acid detection in effusions of cats with suspected feline infectious peritonitis. Res Vet Sci. 2019 Aug;125:421–424. doi:10.1016/j.rvsc.2017.10.004. Epub 2017 Oct 31. PMID: 29113645; PMCID: PMC7111774.

39. Herrewegh AA, de Groot RJ, Cepica A, Egberink HF, Horzinek MC, Rottier PJ. Detection of feline coronavirus RNA in feces, tissues, and body fluids of naturally infected cats by reverse transcriptase PCR. J Clin Microbiol. 1995 Mar;33(3):684–9. doi: 10.1128/jcm.33.3.684-689.1995. PMID: 7751377; PMCID: PMC228014.

40. Dye C, Helps CR, Siddell SG. Evaluation of real-time RT-PCR for the quantification of FCoV shedding in the faeces of domestic cats. J Feline Med Surg. 2008 Apr;10(2):167–74. doi: 10.1016/j.jfms.2007.10.010. Epub 2008 Feb 20. PMID: 18243744; PMCID: PMC2582154.

41. Hirotsu Y, Mochizuki H, Omata M. Double-quencher probes improve detection sensitivity toward Severe Acute Respiratory Syndrome Coronavirus 2 (SARS-CoV-2) in a reverse-transcription polymerase chain reaction (RT-PCR) assay. J Virol Methods. 2020 Oct;284:113926. doi: 10.1016/j.jviromet.2020.113926. Epub 2020 Jul 7. PMID: 32650037; PMCID: PMC7341737.

42. Nunes BTD, de Mendonça MHR, Simith DB, Moraes AF, Cardoso CC, Prazeres ITE, de Aquino AA, Santos Adcm, Queiroz ALN, Rodrigues DSG, Andriolo RB, Travassos da Rosa ES, Martins LC, Vasconcelos PFDC, Medeiros DBA. Development of RT-qPCR and semi-nested RT-PCR assays for molecular diagnosis of hantavirus pulmonary syndrome. PLoS Negl Trop Dis. 2019 Dec 26;13(12):e0007884. doi: 10.1371/journal.pntd.0007884. PMID: 31877142; PMCID: PMC6932758.

43. Bui HTV, Bui HT, Chu SV, Nguyen HT, Nguyen ATV, Truong PT, Dang TTH, Nguyen ATV. Simultaneous real-time PCR detection of nine prevalent sexually transmitted infections using a predesigned double-quenched TaqMan probe panel. PLoS One. 2023 Mar 6;18(3):e0282439. doi: 10.1371/journal.pone.0282439. PMID: 36877694; PMCID: PMC9987813.

44. Gunn-Moore DA, Gruffydd-Jones TJ, Harbour DA. Detection of feline coronaviruses by culture and reverse transcriptase-polymerase chain reaction of blood samples from healthy cats and cats with clinical feline infectious peritonitis. Vet Microbiol. 1998 Jul;62(3):193–205. doi: 10.1016/s0378-1135(98)00210-7. PMID: 9791867; PMCID: PMC7117229.

