## Supplemental Figures for "Detection of Feline Coronavirus RNA in Cats with Feline Infectious Peritonitis and their Housemates"

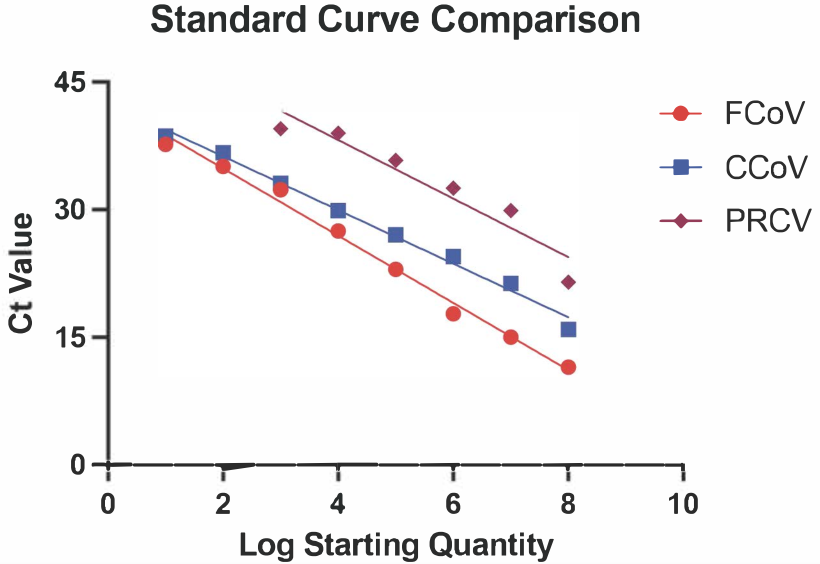


**Supplemental Figure 1: Assay cross-reactivity with similar Alphacoronaviruses.** Standard curves ranging from 1.0E8 to 1.0E1 were generated for three coronaviruses: Feline (FCoV, red), Canine (CCoV, blue), and Porcine (PRCV, purple). Graph displays decreased assay sensativity for CCoV at higher amount of standard, but comparable levels at lower amounts compared to FCoV standards (compare red to blue). Loss of sensitivity is greater when comparing PRCV and FCoV at all amounts tested (compared red to purple).

| | Control Cat 001 | Control Cat 002 | Control Cat 003 | Control Cat 004 | Control Cat 005 | Control Cat 006 | Control Cat 007 | Control Cat 008 | Control Cat 009 |
|---|---|---|---|---|---|---|---|---|---|
| Plasma | Below LoD | Below LoD | Below LoD | Below LoD | Below LoD | Below LoD | Below LoD | NA | Below LoD |
| Whole Blood | Below LoD | Below LoD | 1.7e5 RNA copies/ml | Below LoD | Below LoD | Below LoD | Detected | Below LoD | Below LoD |
| Fecal Swabs | Detected | Below LoD | 2.9e6 RNA copies/swab | Below LoD | Below LoD | 5.6e7 RNA copies/swab | Below LoD | Below LoD | Below LoD |
| Conunctival Swabs | 6.5e6 RNA copies/swab | 6.0e6 RNA copies/swab | Detected | Detected | Detected | Detected | Detected | Detected | Detected |

**Supplemental Table 1: qRT-PCR results from nine unexposed, healthy cats.** Quantifiable samples have numerical values. LoD, limit of detection. NA, not applicable due to low sample availability. Samples testing below the limit of quantification but above the limit of detection are labelled as “Detected”.
